# Estimation of swine movement network at farm level in the US from the Census of Agriculture data

**DOI:** 10.1101/488767

**Authors:** Sifat A. Moon, Tanvir Ferdousi, Adrian Self, Caterina M. Scoglio

## Abstract

Swine movement networks among farms/operations are an important source of information to understand and prevent the spread of diseases, nearly nonexistent in the United States. An understanding of the movement networks can help the policymakers in planning effective disease control measures. The objectives of this work are: 1) estimate swine movement probabilities at the county level from comprehensive anonymous inventory and sales data published by the United States Department of Agriculture - National Agriculture Statistics Service database, 2) develop a network based on those estimated probabilities, and 3) analyze that network using network science metrics. First, we use a probabilistic approach based on the maximum entropy method to estimate the movement probabilities among different swine populations. Then, we create a swine movement network using the estimated probabilities for the counties of the central agricultural district of Iowa. The analysis of this network has found evidence of small-world phenomenon. Our study suggests that the US swine industry may be vulnerable to infectious disease outbreaks because of the small-world structure of its movement network. Our system is easily adaptable to estimate movement networks for other sets of data, farm animal production systems, and geographic regions.

## Introduction

Livestock is often moved between facilities to reduce costs and improve productivity. There is an old adage, “Livestock follow the grain”. Even now this proverb seems true, as shipping the animals is less expensive than shipping the grains, which are required for animals to attain their slaughter weights. The corn-belt region (Iowa, Missouri, Illinois, Indiana, and Ohio) is the largest market for feeder pigs^1^ because they are the largest producers of two major sources of hog rations (corn and soybeans). Although movements in the livestock industry can reduce the cost of production, movements have a major role in the risk of pathogens spread. Movement of swine among the farms is one of the major pathways for the spreading of several diseases (e.g., PRRS-Porcine reproductive and respiratory syndrome, PED-Porcine epidemic diarrhea etc.) in the US swine industry^2,3^. Knowledge of livestock movement could be useful in the control of pathogen spread. In Europe, there are several well-established animal tracking systems. However, similar programs are yet to be mandated for the US. In the US, a comprehensive livestock tracking system has not been implemented because of a cultural preference for privacy and competition between producers^4^. The United State Department of Agriculture collects movement information when livestock shipments cross state boundaries. There is no program that collects movement information at the county or farm level.

In the past literature, several models have been developed to understand swine movement in different regions of the US^4–6^. However, all of them used confidential incomplete datasets, which are not publicly accessible, and also which are not inclusive of the whole US. Yadav et al.^5^ developed a model to understand classical swine fever outbreak-related outcomes in Indiana. They have used data from USHerds (US Animal Health Emergency Reporting and Diagnostic System), where import-export activities, location of import origin, receiving swine premises, shipment size and shipment date are listed. However, only 22% of the states participates in the USHerds program. Another research group predicted movement networks of the swine industry for some counties of Minnesota using a machine learning approach^6^. They used confidential survey data from two counties to train their model. The objective of our research is to understand the swine movement network in the US from publicly available census data. A network is a useful structure in the study of any spreading phenomena, where farm-level animal movement networks are used as a key component in the area of disease spreading^7,8^.

In this work, we estimate the swine movement probabilities between counties based on published inventory and sales data from the Census of Agriculture. We develop a convex optimization problem with some linear constraints for the US swine industry. To solve this problem, we adapt the cattle movement model from Schumm et al.^9^ to the swine population. In particular, we maximize the entropy of out-going distributions from each swine sub-population. Maximum entropy methods have been used in various research fields^10–12^. The maximum entropy principle states that the best way to approximate the unknown distribution that satisfies all the constrains will have the maximum entropy^13^. We propose a novel algorithm to develop a farm level swine movement network using the estimated swine movement probabilities. In this network, nodes (or vertices) represent the swine-farms and the directed links (or edges or connections) represent directional swine movements between the farms. To understand the generated swine movement network, we use network analysis tools. It has been used several times in the literature to understand the livestock movement patterns^14–16^. From the analysis of the developed swine movement network, we find trace of the small world phenomenon, and presence of hubs in the US swine movement network.

## Materials and Methods

First, we develop a convex optimization problem to estimate swine movement probabilities. Next, we propose an algorithm to develop a network based on those probabilities, where nodes or vertices are farms or operations and edges among them represent swine movement. Finally, we analyze the network using different network analysis metrics.

### Data

We collect the county-wise hog inventory, sales, slaughter, and dead/lost pig data from the United States Department of Agriculture National Agricultural Statistics Service (USDA-NASS)^17^. The USDA-NASS conducts a census every five years, which compiles a uniform, comprehensive agricultural data set for each county of the entire US. We used the data from the 2012 Census of Agriculture, as the census of 2017 is not published fully at the time of this research. For each county, two sets of data are available: 1) inventory and 2) sales. In both types, pigs are grouped into seven classes based on operation/farm size. These groups are: size1 (1-24 pigs), size2 (25-49 pigs), size3 (50-99 pigs), size4 (100-199 pigs), size5 (200-499 pigs), size6 (500-999 pigs), and size7 (more than 1000 pigs). For each size group, data for the number of operations and the number of pigs are available. However, several data points were not published to maintain anonymity; we estimate those to develop the network model. Another set of missing data are the geographic farm locations; we use geographical county centroids to measure the distances among counties.

We estimate the swine movement probabilities among sub-populations for the state of Iowa, where a sub-population is denoted as the swine population in a size group in a county. Iowa has the largest swine inventory (31.43%) in the US^18^. In the list of America’s top 100 pig farming counties, 42 counties are from Iowa alone^19^. It is also the most vulnerable state for the introduction of classical swine fever and African swine fever viruses due to legal import of live swine^20^. Having Iowa 99 counties in total, the number of swine sub-populations in our optimization problem is 99 × 7.

### Swine movement probability estimation

To estimate the pig movement rate among different sub-populations, we use a convex optimization problem. This convex optimization problem consists of two steps: 1) estimation of the non-disclosed data points in the inventory and sales data and 2) estimation of movement probabilities among different sub-populations.

To estimate non-disclosed points in the inventory data, we formulate an entropy function. By maximizing this function, we estimate the data points with minimum assumptions^21^. This process is detailed in Schumm et al^9^. In step 2, we construct a convex optimization problem, which includes a series of linear constraints. The objective of this problem is to maximize the entropy of the out-going flows from each sub-population. The maximum entropy is a well-known method of statistical inference, which has been used in diverse research fields including ecology, thermodynamics, economics, forensics, language processing, astronomy, image processing etc.^12,22,23^. This method produces the least biased predictions while maintaining prior knowledge constraints.

In the convex optimization problem, there are *C* counties and each county has *I* size groups. A pig from a sub-population could be moved to other sub-population, or slaughtered, or lost, or stayed in that sub-population. We define the entropy of the out-going distributions from all sub-populations as,

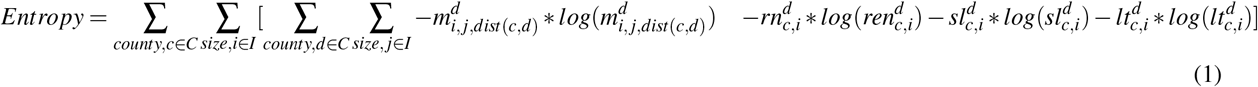

The purpose of this step is to find the decision variables that maximize the Eq 1. We estimate the movement parameter 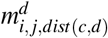, which represents the movement probability from sub-population (*c, i*) to sub-population (*d, j*). A sub-population (*c, i*) is the swine population in the size group *i* in the county *c*. The superscript *d* marks the decision variable. 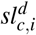 is the probability of pigs being slaughtered for meat from sub-population (*c, i*). 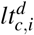 is the probability of pigs being dead or lost in sub-population (*c, i*). 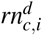 is the probability of remaining in sub-population (*c, i*). We divide the distance between counties into five classes: 1) distance ∈ [0,20), 2) distance ∈ [20,100), 3) distance ∈ [100,200), 4) distance ∈ [200,400), and 5) distance ∈ [400, *D_max_*]. *dist*(*c, d*) represents the distance class for the distance between county *c* and *d*. This problem is subject to several linear constraints, which we construct from statistical rules, movement data, and swine population conservation.

The constraints for the statistical rule are,

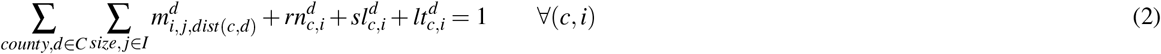

Constraints in the Eq 2 state that the sum of the outgoing probabilities from a sub-population is equal to 1.

Constraints for the movement data are,

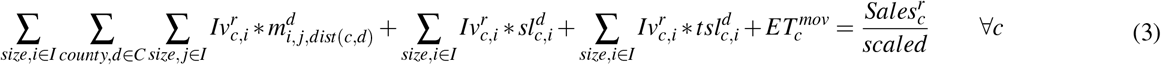

The superscript *r* indicates published data. 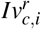 is the swine inventory in sub-population (*c, i*). 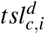 represents sales from sub-population (*c, i*) to outside of the state. 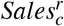 represents the total sales from county *c*. The term *scaled* is used to convert the timescale. The data here are yearly data, this term allows us to convert the timescale from yearly to weekly basis. 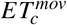 is the error term for movement constrains.

Constrain for the slaughtered swine is,

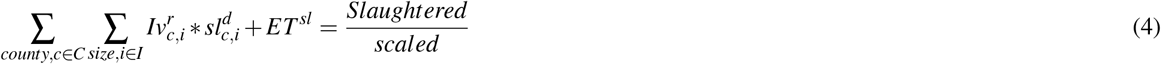

The term *Slaughtered* represents the total number of slaughtered in a year in the system. *ET^sl^* is the error term for slaughtered data.

Constrain for the inshipments is;

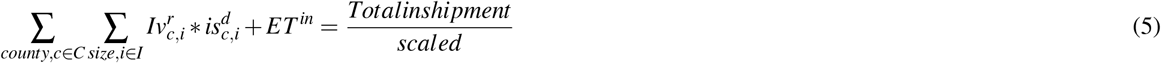

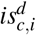 indicates the inshipment rate from outside of the state in sub-population (*c, i*). *Totalinshipment* is the inshipment from outside in a year in the system. *ET^in^* is the error term for inshipment.

In a sub-population, summation of the outgoing flows is equal to the summation of the incoming flows. Constrains for the population conservation are,

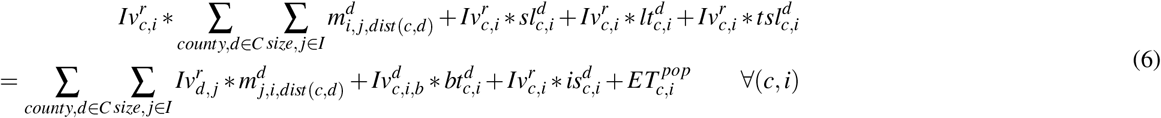

here, 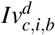 represents the breeding population, 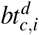 is the probability of birth in county *c* and size group *i*, and 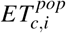 is the error term. The left side of the equation 6 is the summation of the outgoing flows from sub-population (*c, i*) and the right side is the summation of the incoming flows into the sub-population (*c,i*). The range for 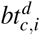 is (7 × 9)/115 − (7 × 12)/112 *week*^−1^, as time period for gestation is 112-115 days and average litter rate is 9–12^18^. The range for 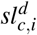 was chosen based on the lifespan of market pigs in the US, which is about 25 to 28 weeks.

Constrain for the errors is,

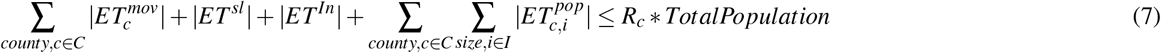

The left side of the Eq 7 represents the summation of all the errors in the optimization problem. Here, *R_c_* is a proportional constant, and *TotalPopulation* is the total swine population in the system.

Convex cost function (Eq 1) and constrains (Eq 2–7) constitute our optimization linear problem.

### Network Development

We develop a network using the movement parameters which are obtained using the maximum entropy optimization. The network development is done in two stages: 1) setup of the population in each farm and 2) setup of the movement links between farms.

In order to generate the network, first, we need the farm level estimates of the pig population. The USDA-NASS data only provide the number of farms in a size range and the number of total pigs in that range. Recorded data on the number of pigs in a farm are generally not available in the US (with the exception of a few counties). To allocate the pig population, we generate random numbers for every farm in a size group *i* within a county *c* with the following constraints:

a. The random numbers fall in the range of the corresponding group *i*.
b. The sum of all generated numbers is equal to the total number of pigs in that sub-population (*c, i*).

Our movement network for pig farms is represented as (*V, E, W*). The term *V* denotes the set of nodes, the term *E* represents the set of links or connections among individual nodes, and *W* denotes the weight of each link. To generate the movement network among farms, we use the following procedures:

Step 1 For each pig *p*_1_ in a sub-population (*c, i*), we generate a random number *rand* for sub-population (*d, j*), *d* = 1,2,3,*C*, and *j* = 1,2,3, ….*I*. Here, *C* is the number of counties in the system and *I* is the number of size groups.
Step 2 If 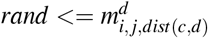, a link is created from pig *p*_1_ to another pig *p*_2_. Pig *p*_2_ is picked randomly from the sub-population (*d, j*).
Step 3 If there is no link from the parent farm *f*_1_ of pig *p*_1_ to the parent farm *f*_2_ of pig *p*_2_, we create a link *flink* from *f*_1_ to *f*_2_. Otherwise, if a link already exists, we increase its weight by 1.
Step 4 For each sub-population (*c, i*), we repeat Steps 1-3.

This process produces a directed weighted network at the farm level. Links or connections among farms represent swine movement. The weight of a link represents the volume of movements occurred from one farm to another.

### Network Analysis

To capture the particular features of the developed network, we compute the following network analysis metrics: node strength, betweenness, eigenvector, clustering coefficient centrality measures, and average shortest path^24–26^. Centrality measures can help us determine the most important or central nodes in a network.

The **node strength-centrality** measure is the strength of the nodes or sum of the weights of the edges connected to it^27^. In a directed network, the nodes have two types of vertex-strength centralities: 1) in-strength *Is*, and 2) out-strength *Os*.

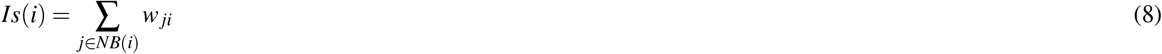

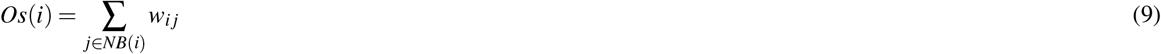

Here, *w_ji_* is the connection strength of the edge/link from *j* to *i, NB*(*i*) is the set of the neighbors of node *i*. Vertex strength can be illuminating in the investigation of diseases spreading. A high in-strength node has a high risk of receiving infection. On the other hand, a high out-strength node is influential over the network, as such a node can infect many more nodes.

The **betweenness centrality** measure suggests which nodes are important in the connection flow or act as bridges in the network. Betweenness centrality of a node measures how many shortest path between different pairs of nodes go through that particular node. Nodes with high betweenness centrality have high control over flow (here, concerning flow of swine) in the network. Removal of such nodes can effectively reduce connectivity in the network. Knowledge of these nodes can be useful in controlling outbreaks^28^. Let, *p_st_* be the number of shortest paths from *s* ∈ *N* to *t* ∈ *N*. We denote, *p_st_*(*i*) to be the number of shortest paths from *s* to *t*, that includes node *i* somewhere in between. The betweenness centrality of a node *i* is defined^29^ as:

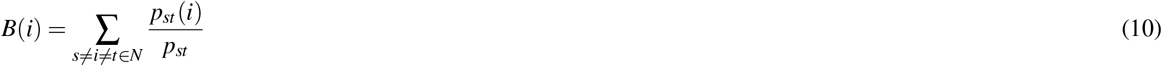

**Eigenvector centrality** is an extension of the degree/strength centrality. In the eigenvector centrality measure, the centrality of a node is proportional to the sum of the centralities of its neighbors.

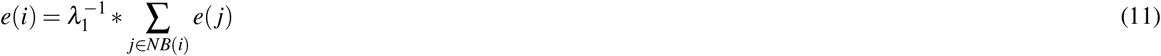

Here, *e*(*i*) is the eigenvector centrality of the node *i*, and *λ*_1_ is the largest eigenvalue of the adjacency matrix [*a_ij_*] of the network. Eigenvector centrality of a node can be large if either it has many neighbors or it has important neighbors. Nodes with high eigenvector centralities have high probabilities of becoming infected^30,31^.

The **clustering coefficient** measures local group cohesiveness. The clustering coefficient *Cc*(*i*) for a node *i* is the ratio of the number of edges among the neighbors of *i* and the maximum possible number of such edges (for the fully-connected network formed by the neighbors of node *i*). If neighboring nodes of node *i* has *c_i_* connections among them then clustering coefficient can be defined as^25^:

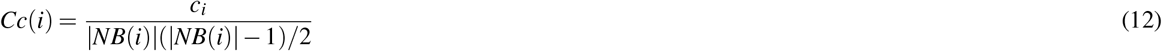

## Results

### Movement probability estimation

In this research, we solve a convex optimization problem to estimate the swine movement probabilities by using the maximum entropy approach for Iowa. We utilized the AIMMS modeling system^32^ of Paragon Decision Technology to solve this convex optimization problem. One assumption of this problem is sub-population sizes are constant. The time-scale of our estimation problem is weekly, which we controlled it by using *scaled* = 52*weeks/year*. The boundary of error limit in our system is *R_c_* = 5.5% of total swine population in Iowa. We found this value from trial and error with an objective to minimize the total error. The estimated probabilities are given in Table 1. Table 1 is showing swine movement probabilities between size groups for five different distance ranges. The highest movement probability is from size7 to size7 sub-population, when the distance between them is less than 20km. We divide seven size groups into three categories; size: 1-3(small farms), 4-5(medium farms), and 6-7(large farms). The probability is small for the movement from large farms to small farms and vice versa.

**Table 1.** Estimated swine movement probabilities *m_i,j,dist(c,d)_* × 10^3^ from maximum entropy approach.

|  |  | Destination |  |  |  |  |  |  |
| --- | --- | --- | --- | --- | --- | --- | --- | --- |
|  |  | Size1 | Size2 | Size3 | Size4 | Size5 | Size6 | Size7 |
| **Distance < 20km** |  |  |  |  |  |  |  |  |
| Source | size1 | 1.4959 | 1.3737 | 1.4112 | 1.4056 | 1.4410 | 1.4556 | 1.4911 |
|  | size2 | 1.4333 | 1.4933 | 1.4073 | 1.4163 | 1.4350 | 1.4444 | 1.4951 |
|  | size3 | 1.3647 | 1.2582 | 1.6877 | 1.4917 | 1.5190 | 1.5384 | 1.5859 |
|  | size4 | 1.2084 | 1.2435 | 1.3374 | 2.0952 | 1.7075 | 1.7960 | 1.9354 |
|  | size5 | 0.2962 | 0 | 0.8019 | 1.9487 | 5.7582 | 4.9944 | 4.6968 |
|  | size6 | 0 | 0 | 0 | 1.4524 | 7.6968 | 10.9211 | 7.8967 |
|  | size7 | 0 | 0 | 0 | 0 | 0 | 0.8257 | 54.2762 |
| **20km < Distance < 100km** |  |  |  |  |  |  |  |  |
| Source | size1 | 1.3813 | 1.3312 | 1.3876 | 1.4055 | 1.4352 | 1.4441 | 1.4858 |
|  | size2 | 1.3807 | 1.3260 | 1.3823 | 1.4070 | 1.4332 | 1.4404 | 1.4843 |
|  | size3 | 1.3236 | 1.2157 | 1.3588 | 1.3865 | 1.4976 | 1.5229 | 1.5861 |
|  | size4 | 1.2212 | 0.9263 | 1.2939 | 1.3162 | 1.6023 | 1.6535 | 1.8475 |
|  | size5 | 0 | 0 | 0.0671 | 0.6336 | 2.4507 | 3.1341 | 4.0366 |
|  | size6 | 0 | 0 | 0 | 0 | 2.1857 | 3.6179 | 5.5171 |
|  | size7 | 0 | 0 | 0 | 0 | 0 | 0 | 0 |
| **100km < Distance < 200km** |  |  |  |  |  |  |  |  |
| Source | size1 | 1.3728 | 1.3205 | 1.3865 | 1.3955 | 1.4400 | 1.4432 | 1.4900 |
|  | size2 | 1.3779 | 1.3278 | 1.3917 | 1.3969 | 1.4360 | 1.4397 | 1.4858 |
|  | size3 | 1.3392 | 1.2196 | 1.3511 | 1.3860 | 1.4725 | 1.4846 | 1.5895 |
|  | size4 | 1.2067 | 0.9283 | 1.2853 | 1.3543 | 1.5697 | 1.6161 | 1.8692 |
|  | size5 | 0 | 0 | 0.22110 | 0.62150 | 2.1443 | 2.3515 | 4.1797 |
|  | size6 | 0 | 0 | 0 | 0 | 1.2824 | 1.7695 | 6.0746 |
|  | size7 | 0 | 0 | 0 | 0 | 0 | 0 | 0 |
| **200km < Distance < 400km** |  |  |  |  |  |  |  |  |
| Source | size1 | 1.3641 | 1.3269 | 1.3851 | 1.3973 | 1.4346 | 1.4487 | 1.4956 |
|  | size2 | 1.3631 | 1.3270 | 1.3846 | 1.3968 | 1.4367 | 1.4504 | 1.4935 |
|  | size3 | 1.3090 | 1.2202 | 1.3545 | 1.3799 | 1.4648 | 1.4889 | 1.6106 |
|  | size4 | 1.1813 | 0.9759 | 1.2754 | 1.3477 | 1.5448 | 1.5977 | 1.8757 |
|  | size5 | 0 | 0 | 0.1522 | 0.6706 | 1.9323 | 2.2975 | 4.4027 |
|  | size6 | 0 | 0 | 0 | 0 | 0.6692 | 1.4190 | 6.7712 |
|  | size7 | 0 | 0 | 0 | 0 | 0 | 0 | 0 |
| **Distance > 400km** |  |  |  |  |  |  |  |  |
| Source | size1 | 1.3175 | 1.3092 | 1.3687 | 1.4282 | 1.4343 | 1.4829 | 1.5128 |
|  | size2 | 1.3434 | 1.3050 | 1.3726 | 1.4186 | 1.4300 | 1.4783 | 1.5049 |
|  | size3 | 1.2390 | 1.1926 | 1.3274 | 1.4639 | 1.4350 | 1.5919 | 1.6360 |
|  | size4 | 1.1469 | 0.8761 | 1.3004 | 1.4588 | 1.4765 | 1.7712 | 1.8739 |
|  | size5 | 0 | 0 | 0 | 0.6439 | 1.2650 | 3.5239 | 4.1429 |
|  | size6 | 0 | 0 | 0 | 0 | 0 | 3.2889 | 5.6178 |
|  | size7 | 0 | 0 | 0 | 0 | 0 | 0.0004 | 0.0029 |

### Network description

We generate a swine movement network for the central agricultural district of Iowa. It has 12 counties: Boone, Dallas, Grundy, Hamilton, Hardin, Jasper, Marshall, Polk, Poweshiek, Story, Tama, and Webster. The total number of farms in those 12 counties is 641, while the net pig population is 2,600,888, which is 12.71% of the total pig population in Iowa. Grundy, Hamilton, Hardin, Jasper, Marshall, and Webster County are within the America’s top 100 pork producer counties. Among these, Hardin County is in the 9^th^ position. The descriptions of pig inventories for the above-mentioned counties are provided in the supplementary material Dataset 1.

For these 12 counties, we have developed a movement network (*V,E,W*), which is shown in Fig 1. This network is a realization based on the movement probabilities from Table 1. For the network, |*V*| = 641 and |*E*| = 26,060, the description of the nodes, and the adjacency list for this network is provided in the supplementary material Dataset 2 and 3. In Fig 1, this network has seven types of nodes representing the seven size groups. A description of size groups are presented in Table 2. The largest group is the size7, contains 393 nodes which are presented by light blue. There are 17484 edges among the the nodes of this group (67.41% of total edges).

**Figure 1.**
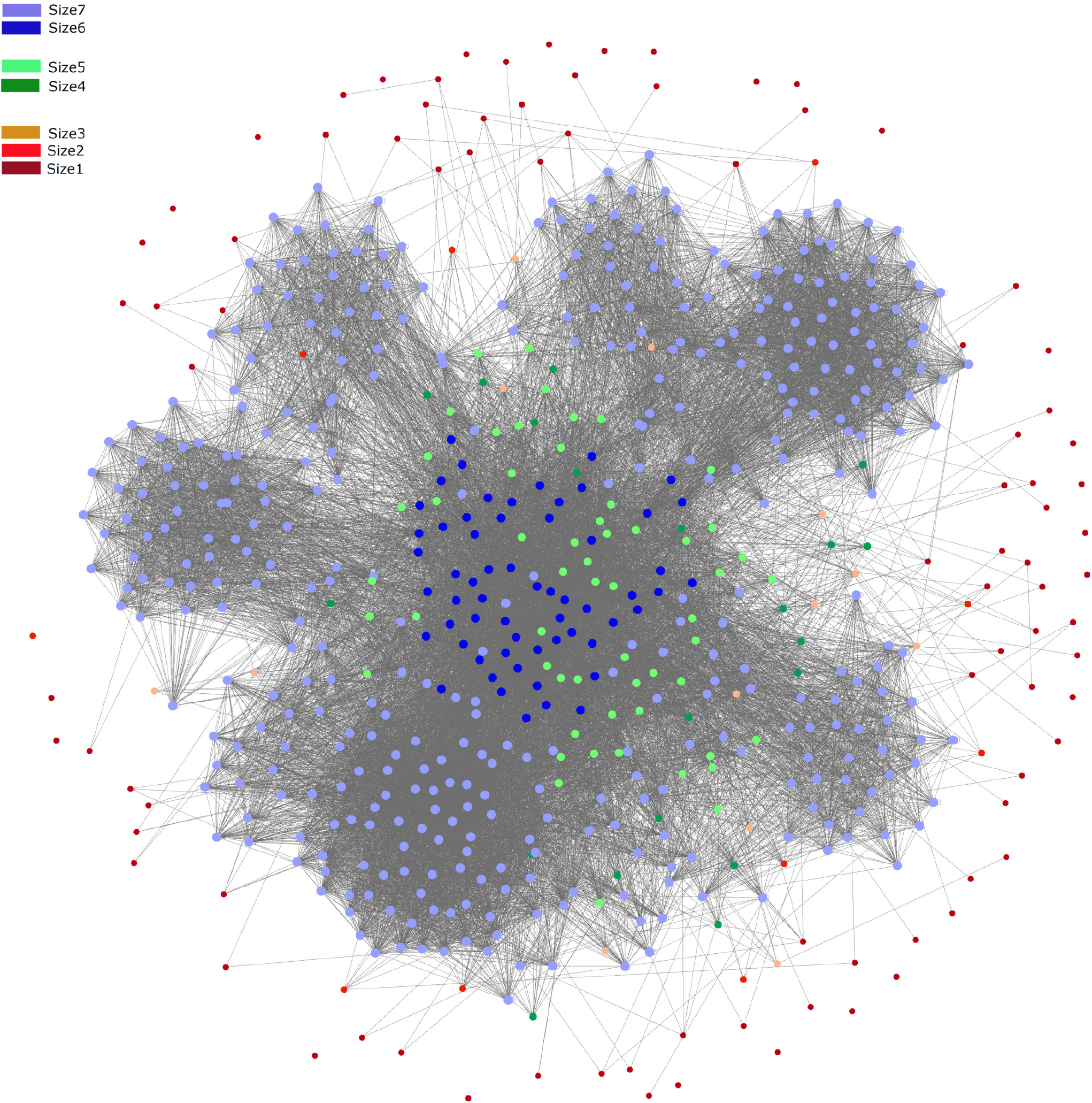
Movement Network for the pig population in the farm level. Different color represents different size groups. Farms are divided into 7 size groups, size: 1-3(small farms), 4-5(medium farms), and 6-7(large farms).

**Table 2.**
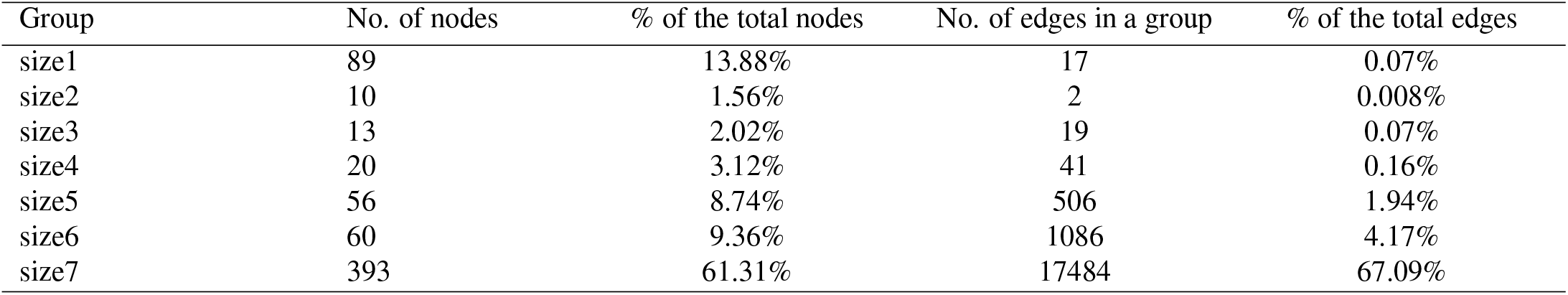
A summary of the size groups in the network.

| Group | No. of nodes | % of the total nodes | No. of edges in a group | % of the total edges |
| --- | --- | --- | --- | --- |
| size1 | 89 | 13.88% | 17 | 0.07% |
| size2 | 10 | 1.56% | 2 | 0.008% |
| size3 | 13 | 2.02% | 19 | 0.07% |
| size4 | 20 | 3.12% | 41 | 0.16% |
| size5 | 56 | 8.74% | 506 | 1.94% |
| size6 | 60 | 9.36% | 1086 | 4.17% |
| size7 | 393 | 61.31% | 17484 | 67.09% |

### Network Analysis

The clustering coefficient of the full network is 0.417, the diameter of the network is 8, and the average shortest path length is 2.725. A summary of various centrality measures for the network is provided in Table 3. Node-strength, betweenness, eigenvector and clustering coefficient centrality for seven size groups are presenting here. In-strength, out-strength, betweenness, and eigenvector centralities were calculated from the overall network. Clustering coefficients in Table 3 were calculated for networks of the same size group (any node and its neighbors are in the same size group). We used the open source package Gephi to visualize and analyze the network^33^.

**Table 3.** A summary of centrality measures for different size groups in the network.

|  | Size1 | Size2 | Size3 | Size4 | Size5 | Size6 | Size7 |
| --- | --- | --- | --- | --- | --- | --- | --- |
| **In-strength** |  |  |  |  |  |  |  |
| mean | 1.202 | 6.300 | 9.846 | 15.250 | 38.232 | 79.117 | 417.791 |
| median | 1.000 | 4.500 | 8.000 | 12.000 | 21.000 | 32.000 | 262.000 |
| (95% CI) | (0.763,<br>1.641) | (1.174,<br>11.426) | (5.232,<br>14.461) | (8.192,<br>22.309) | (21.235,<br>55.229) | (51.804,<br>106.430) | (375.080,<br>460.503) |
| range | (0, 11) | (0, 24) | (1, 29) | (7, 76) | (7, 451) | (14, 543) | (41, 2633) |
| **Out-strength** |  |  |  |  |  |  |  |
| mean | 1.067 | 3.500 | 11.308 | 18.750 | 50.464 | 124.333 | 409.020 |
| median | 1.000 | 3.500 | 10.000 | 17.000 | 44.500 | 104.5000 | 270.000 |
| (95% CI) | (0.760,<br>1.375 ) | (1.772,<br>5.228) | (6.408,<br>16.207) | (14.251,<br>23.249) | (44.010,<br>56.919) | (110.147,<br>138.520) | (370.854,<br>447.187) |
| range | (0, 7) | (0, 8) | (3, 32) | (7, 39) | (23, 117) | (48, 295) | (49, 2187) |
| **Betweenness** |  |  |  |  |  |  |  |
| mean | 59.580 | 478.316 | 874.537 | 1377.900 | 1329.300 | 5123.400 | 458.014 |
| median | 0 | 338.046 | 600.790 | 1334.900 | 1017.500 | 2606.100 | 325.821 |
| (95% CI) | (23.367,<br>95.790) | (136.166,<br>820.467) | (449.100,<br>1299.900) | (1021.400,<br>1734.500) | (1041.400,<br>1617.300) | (3539.800,<br>6707.100) | (411.019,<br>505.011) |
| range | (0, 737.345) | (0, 1195.4) | (91.2106,<br>2279.3) | (141.363,<br>3391.5) | (65.760,<br>4995.5) | (833.733,<br>36326) | (0.747,<br>4207.9) |
| **Eigenvector** |  |  |  |  |  |  |  |
| mean | .00076 | 0.0066 | 0.0108 | 0.0273 | 0.0794 | 0.1858 | 0.3530 |
| median | .00018 | 0.0036 | 0.0074 | 0.0217 | 0.0662 | 0.0922 | 0.2125 |
| (95% CI) | (0.0005,<br>0.0010) | (0.0003,<br>0.0135) | (0.0045,<br>0.0172) | (0.0177,<br>0.0369) | (0.0641,<br>0.0946) | (0.1289,<br>0.2427) | (0.3257,<br>0.3804) |
| range | (0, 0.0050) | (0, 0.0324) | (.00098,<br>0.0371) | (0.0074,<br>0.1018) | (0.0191,<br>0.3039) | ( 0.0407, 1) | (0.0417,<br>0.9760) |
| **Clustering coefficient** |  |  |  |  |  |  |  |
| mean | 0.006 | 0 | 0.187 | 0.112 | 0.223 | 0.354 | 0.796 |

From the node-strength centrality measures, we observe that the average node-strength is positively correlated with the size groups. Larger size groups have higher average node-strengths. Consequently, size7 has the highest average node-strength (Table 3). The node-strength distribution is provided in Fig 2. In the network, only a few nodes have high strength and most of the nodes have low strength. This characteristic is similar to the power-law distribution. The range of in-strength is 0 – 2633. About 84.24% of the total nodes have in-strengths less than 526, which is merely the first 20% of the in-strength range. The range for out-strength is 0 – 2187. About 79.88% of the total nodes have out-strengths less than 437, which is within first 20% of the range of out-strength values. The correlation coefficient between in-strength and out-strength is 0.7999, which is an indication of strong correlation.

**Figure 2.**
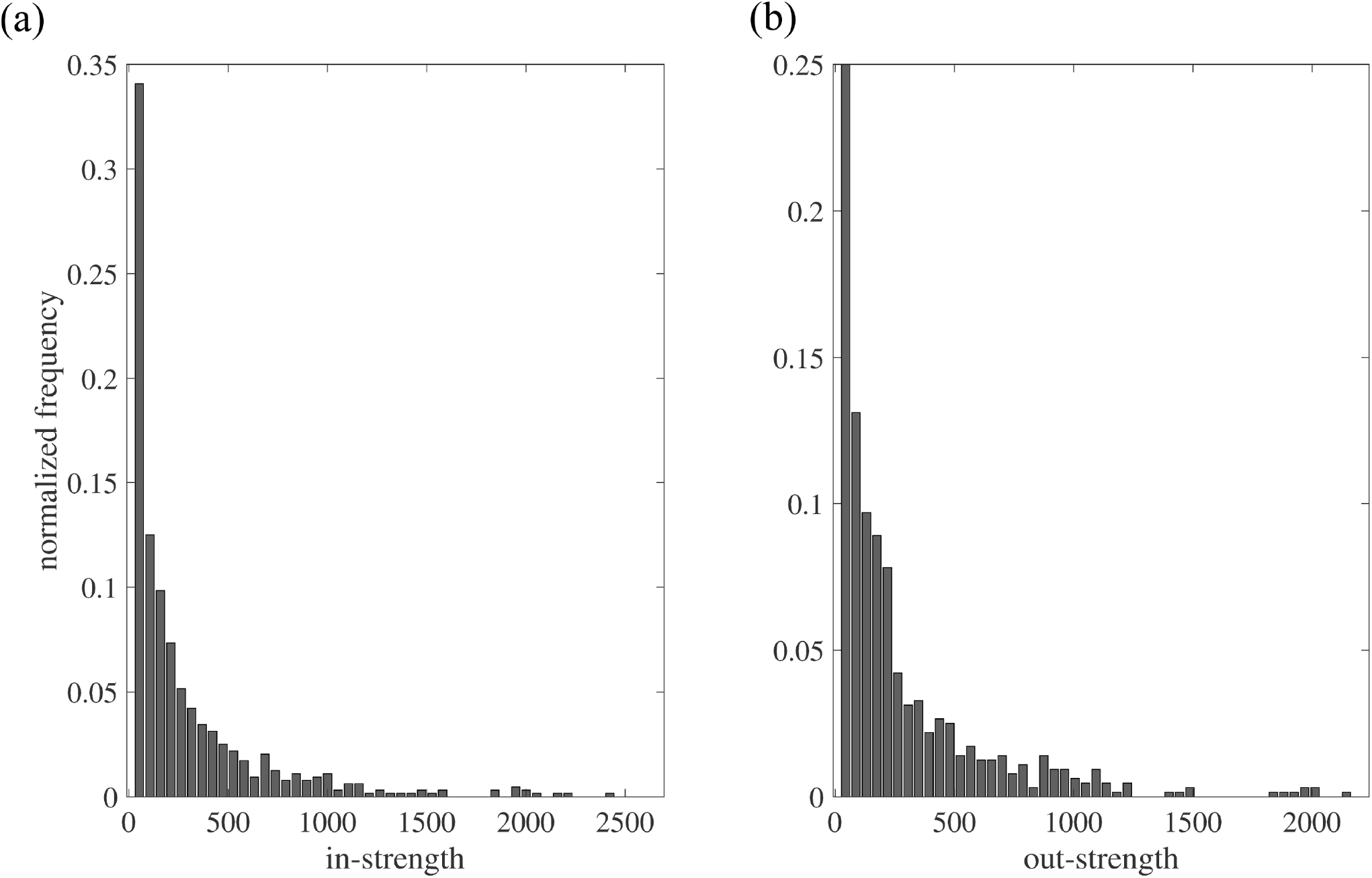
Node strength distribution of the directed network. (a) in-strength, (b) out-strength

The betweenness centrality is positively correlated with size groups until group6, after which farms in the group7 has low betweenness. The farms in group6 have the highest average betweenness. The distribution of betweenness centrality measure is given in Fig 3. Most of the farms have low betweenness. Few farms act as hubs in the network which have high betweenness. We divide the nodes into three groups, 1) low-betweenness (0-100), 2) medium-betweenness (101-1000), and 3) high-betweenness (> 1000). These three groups contain 165, 335, and 141 nodes respectively. These three groups are illustrated in Fig 4. In the low-betweenness group maximum nodes are from small size groups, in the medium-betweenness group most of the nodes are from group7, and in the high-betweenness group, most of the nodes are from group6.

**Figure 3.**
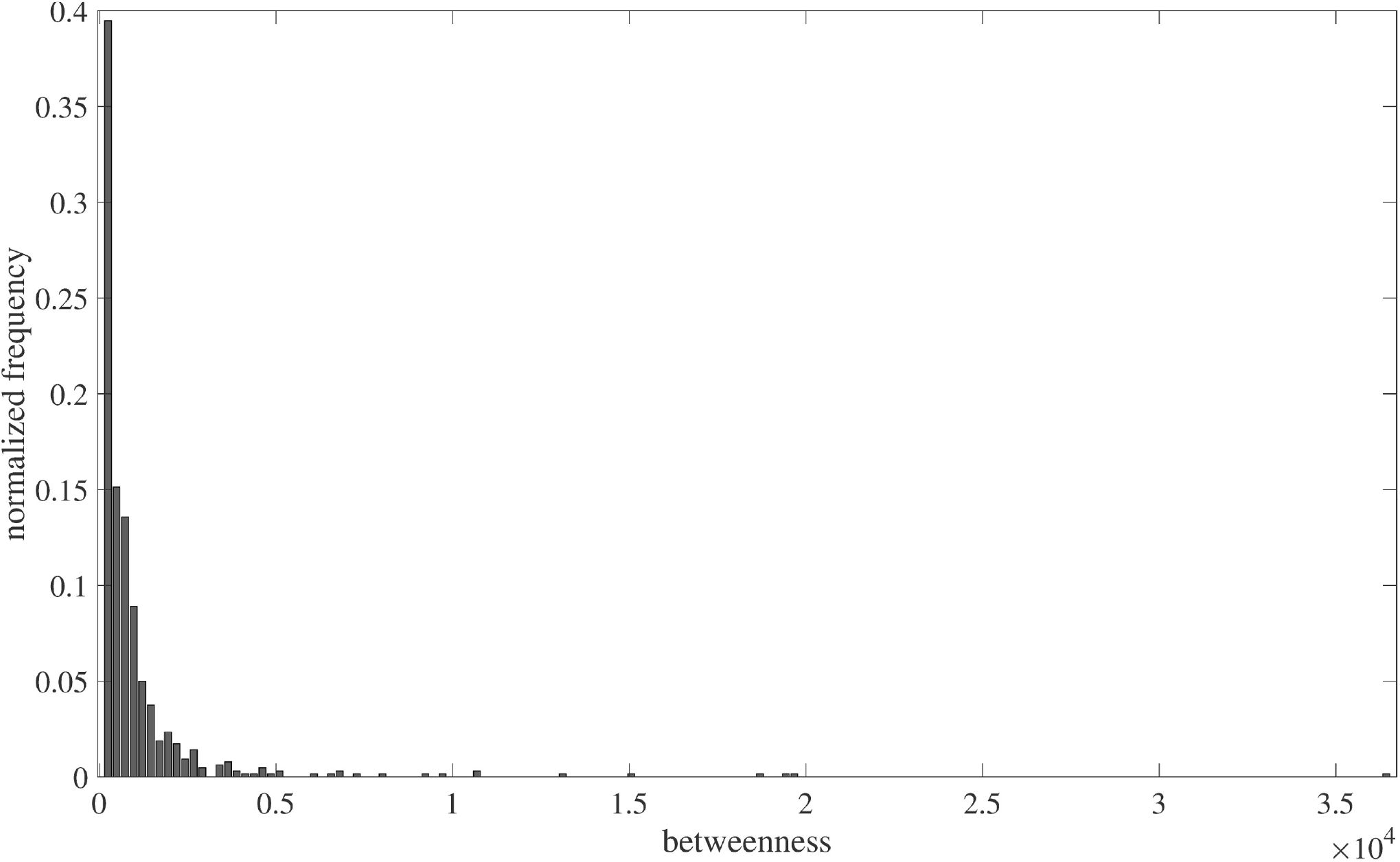
Betweenness distribution of the network.

**Figure 4.**
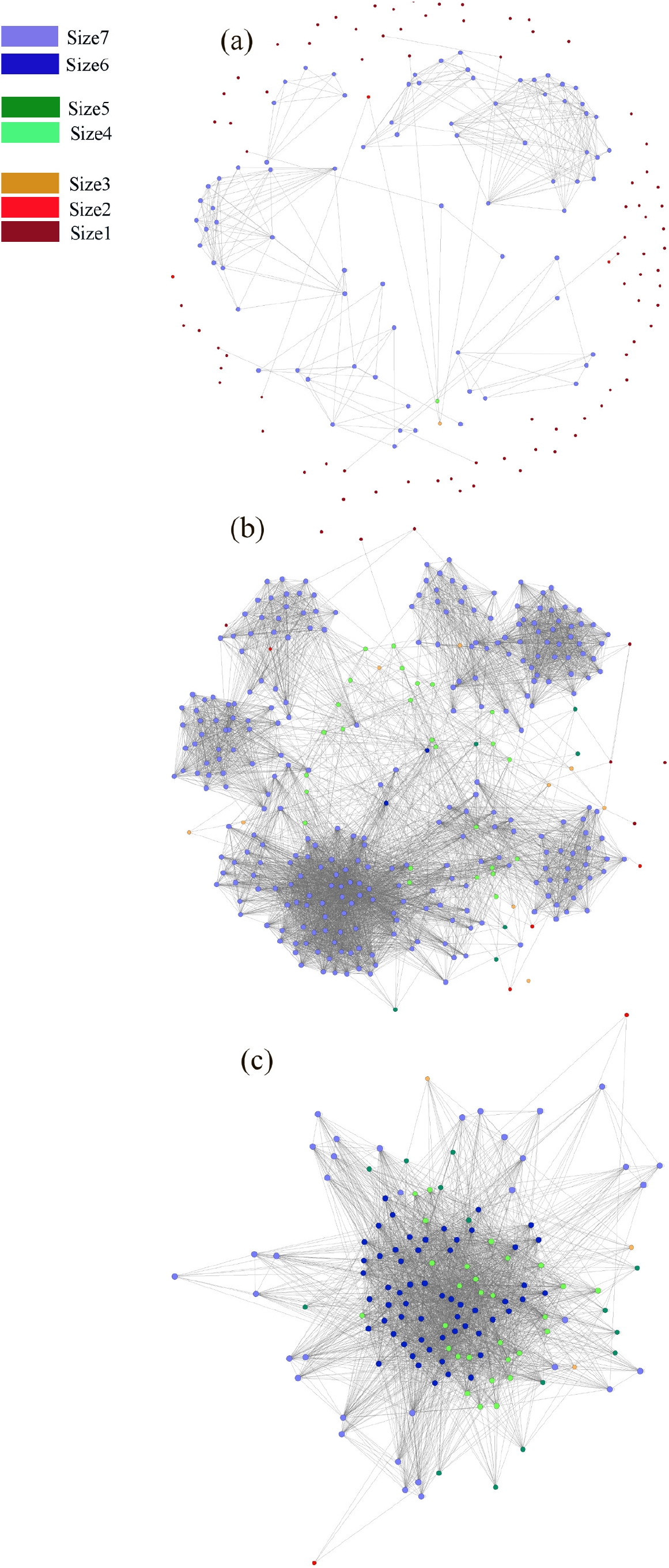
Node groups according to betweenness. a) nodes with low-betweenness, b) nodes with medium-betweenness, and c) nodes with high-betweenness. The connections among visible nodes are presented here.

The mean eigenvector centrality is positively correlated with the size groups. Larger size groups have higher eigenvector centralities (Table 3). We have divided the nodes (farms) into three groups: 1) low-eigenvector central nodes (0-0.1), 2) medium-eigenvector central nodes (0.11-0.5), and 3) high-eigenvector central nodes (0.51-1). The low-eigenvector central group consists of 243 nodes, the medium group consists of 270 nodes, and the high group contains the rest of the nodes. The network for different eigenvector groups is presented in Fig 5. Clustering coefficient for group size 7 is 0.796, which is quite high. The nodes from this group forms several cluster, which are quite visible in Fig 1 and Fig 5.

**Figure 5.**
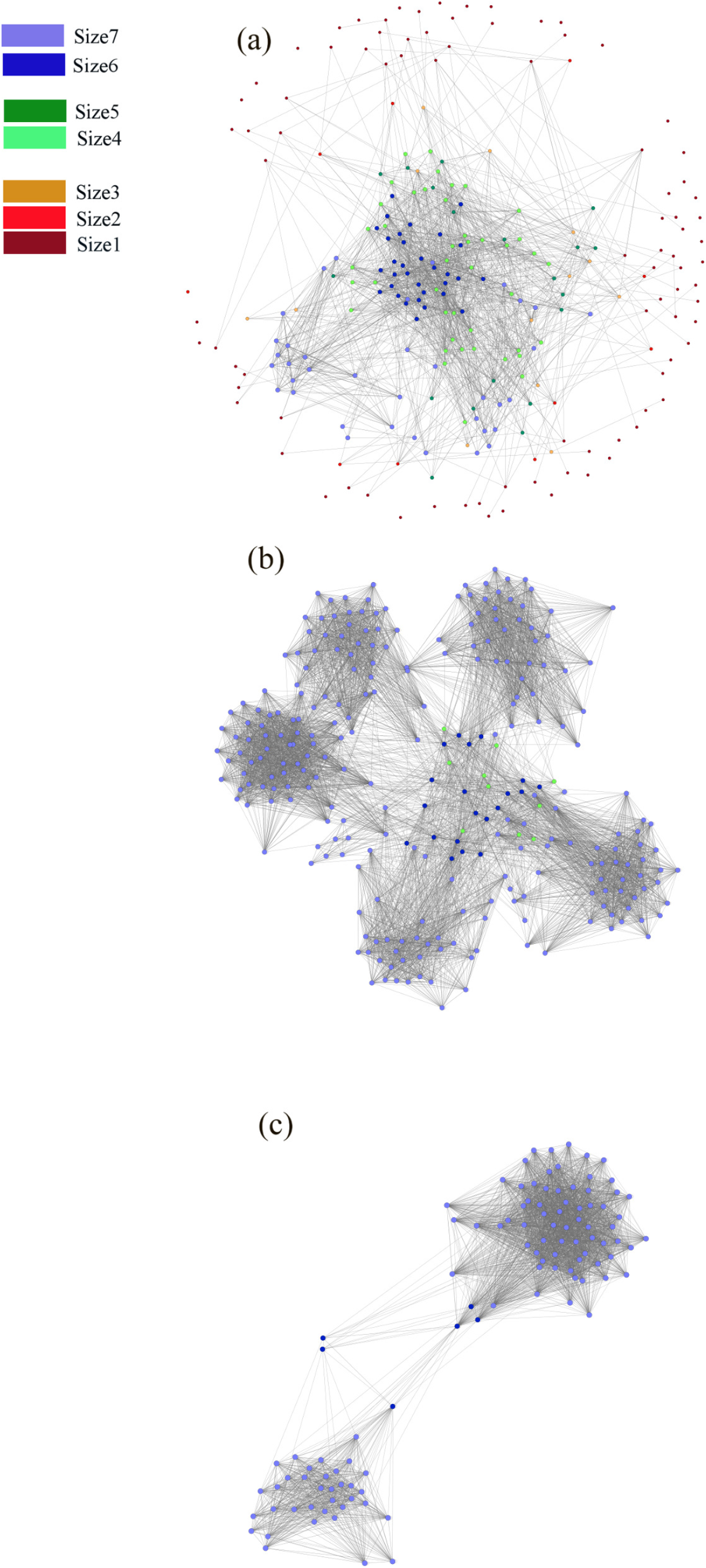
Node groups according to eigenvector centrality. a) Low-eigenvector central nodes, b) medium-eigenvector central nodes, and c) high-eigenvector central nodes. The connections among visible nodes are presented here.

## Discussion

In this study, we have three objectives: 1) we compute optimal estimates swine movement probabilities among counties from the aggregated data of USDA-NASS, 2) we develop a realization of the network from the estimated probabilities, and 3) we analyze the developed network with different network analysis metrics.

Animal movement has been one of the major causes of diseases spread among the farms for several outbreaks in the US swine industry. A better understanding of the swine movement network can increase the feasibility of planning effective mitigation strategies that can reduce the chances of disease spread. There is no mandatory animal movement tracking system in the US due to the appreciation of secrecy in the swine business. We estimated the movements among different swine sub-populations using a convex optimization problem, formulated according to the NASS data. The discrepancy from our optimization problem is about 5.5% of the total swine population, which is slightly higher than that of a similar work on cattle movement probability estimation^9^ due to a greater amount of data available for cattle. Our estimation problem can be improved if more data are available. The information that we have missed the most is the type of pig farms (for example, nursery, farrow-to-feeder, farrow-to-wean, farrow-to-finish, finish only etc.) at the county level. The USDA-NASS department should collect and publish this information in future reports, since this additional data would not hamper the anonymity of the Census of Agriculture but greatly improve movement estimations.

The network development algorithm can provide us a realization of the network from the estimated movement probabilities. The generated swine movement network was well connected with a giant component that contains 95.94% of the farms. Because of this high connectivity, the swine industry is vulnerable to infectious diseases. All the disconnected farms were smaller farms (inventory size less than 100) where most of them produce meat for their own consumption (60.5% of all small swine farms)^34^. In addition to that, most of these small farms are engaged in all of the phases of swine production (farrow-to-finish producers)^35^. On the other hand, larger farms have more connections among them. One possible reason could be that most of the large farms are specialized in a single production phase to increase productivity^36,37^. Consequently, pig shipments are very frequent among them.

We use some centrality measures to understand the characteristics of the movement network. The node-strength distribution of the network is similar to that of scale-free networks (Fig 2). Compared to a random network, epidemics can spread faster in a scale-free network. In addition to that, scale-free networks have lower epidemic threshold transmission rates than comparable random networks^38^. This information could be useful because targeted vaccination/node-removal is more effective in scale-free structures than random vaccination^39^.

If we analyze the average shortest path length and the clustering coefficient of the overall network, we see evidence of the small-world phenomenon in the network. The average path length was much smaller and clustering coefficient was more than six times larger compared to the similar properties of the equivalent Erdos-Renyi random network^40^, which satisfy the sufficient conditions for small-world properties of the network^41^. The US swine movement network structure is quite vulnerable to any pathogen spreading because of its small-world nature. This result is similar to other studies as well^14-16^. This network has high local clustering. Size7 group (larger operations: headcount is more than 1000) has the highest amount of local clustering. Therefore, the large operations are highly interconnected, making them more vulnerable to outbreaks. Moreover, the structure of the US swine industry has been changing over several years. The number of large operations is increasing, where most of them specialize in one particular phase of production. These changes are increasing the risk for disease outbreaks in the swine industry.

The correlation between in-strength (incoming movements) and out-strength (outgoing movements) is strong. The nodes with high out-strength values also have high in-strength values. This is an important indicator as the nodes with high risk of receiving infection are also highly capable of spreading them.

Although the group size7 (largest operations) has the highest values of node-strength, clustering coefficient, and eigenvector centralities it is not necessarily highest in terms of the betweenness centrality measure. We found that the group size6 has the highest betweenness centrality values. The group size4 and size5 also show high betweenness. The above-mentioned properties indicate that the group size7 forms various clusters in the network, where the operations are highly connected. The operations of medium size, however, maintain the connectivity among the clusters of the largest group. Hence, these medium size operations play a key role in the system.

We made several assumptions to simplify our model as all the necessary data were not available. We assumed that the inventory size of the operations is constant on a year-to-year basis. Our network only considers direct movements but there are many indirect ways a disease could spread. Our estimation steps can be easily adapted by adding more constraints when more data are available.

One immediate use of this network could be the investigation of the stochastic spreading processes^42–46^. This kind of study can help us understand the underlying mechanisms and threshold conditions of spreading processes for various swine diseases including porcine reproductive and respiratory syndrome (PRRS), classical swine fever (CSF), African swine fever (ASF) and many more.

In summary, we present a maximum entropy approach to estimate the swine movement network from aggregated anonymous census data. This method can be used to estimate movement probabilities of other farm animals too for various locations.

### Data Availability

The dataset used to perform this research is available from https://quickstats.nass.usda.gov/. The authors are willing to provide further details upon request.

## Supporting information

## Acknowledgements

The authors would like to express their gratitude to Dr. Sanderson for helpful insights into the US swine industry. This material is based upon work supported by the United States Department of Agriculture under the Grant No. 2015-67013-23818 (NIFA) and by the State of Kansas, National Bio and Agro-Defense Facility (NBAF) Transition Fund through the National Agricultural Biosecurity Center (NABC) at Kansas State University.

## Author contributions

S.M., T.F. and C.S. conceived and designed the study, S.M. performed the experiments, S.M. and C.S. analysed the results. S.M. and C.S. wrote the manuscript. T.F., A.S. and C.S. edited the manuscript. All authors reviewed the manuscript.

## Additional information

Competing interests: The authors declare no competing interests.

## Electronic supplementary material

**Dataset 1.** A description of the pig population in the counties of the central agricultural district of IOWA.

**Dataset 2.** A description of the nodes in the network.

**Dataset 3.** Adjacency list.

